# Dual phenotype of MDA-MB-468 cancer cells reveals mutual regulation of tensin3 and adhesion plasticity

**DOI:** 10.1101/130302

**Authors:** Astrid Veβ, Ulrich Blache, Laura Leitner, Angela R.M. Kurz, Anja Ehrenpfordt, Michael Sixt, Guido Posern

## Abstract

Plasticity between adhesive and less-adhesive states is important for mammalian cell behaviour. To investigate adhesion plasticity, we have selected a stable isogenic subpopulation of MDA-MB-468 breast carcinoma cells which grows in suspension. These suspension cells are unable to re-adhere to various matrices or to contract three-dimensional collagen lattices. By transcriptome analysis, we identified the focal adhesion protein tensin3 (Tns3) as a determinant of adhesion plasticity. Tns3 is strongly reduced on mRNA and protein level in suspension cells. Furthermore, challenging breast cancer cells transiently with non-adherent conditions markedly reduces Tns3 expression, which is regained upon re-adhesion. Stable knockdown of Tns3 in parental cells results in defective adhesion, spreading and migration. Tns3 knockdown cells display impaired structure and dynamics of focal adhesion complexes as determined by immunostaining. Restoration of Tns3 expression in suspension cells partially rescues adhesion and focal contact composition. Our work identifies Tns3 as a critical focal adhesion component regulated by, and functionally contributing to, the switch between adhesive and non-adhesive states in MDA-MB-468 cancer cells.

**Summary statement:** We identify the cell-matrix adapter protein tensin3 as a determinant of adhesion plasticity, using cancer cells selected for non-adherent growth. Tensin3 expression constitutes a feedback loop controlling adhesion and motility.

## Introduction

Changing the adhesive properties of cells is important for many biological processes. These include early steps in development like primordial germ cell migration, but also processes in adult organisms such as leukocyte extravasation, dendritic cell homing, neuroregeneration by glia cells, and metastasis of tumor cells (Paluch et al., 2016). In cancer, breakout of cells from solid tumors requires increased motility and reduced cell adhesion. Following their systemic spread via the circulation, the opposite is necessary for the colonization of distal organs, which depends on a (partial) reversion into less motile cells with regained matrix adhesion (Gunasinghe et al., 2012; Chaffer et al., 2016). However, little is known about the cellular regulation of the underlying adhesion plasticity.

Cell-matrix adhesions are macromolecular structures that link the intracellular cytoskeleton to the extracellular matrix (ECM) and are involved in cell attachment, spreading and migration. These adhesion contacts contain the transmembrane integrins, which connect the ECM to the intracellular actin cytoskeleton via adapter proteins (Sun et al., 2016). It is evident that the precise molecular composition of these integrin-based adhesion complexes is critical for their function. Depending on the cell type the so-called integrin “adhesome” can vary a lot and several hundreds of proteins contribute to the “adhesome” as revealed by proteomic analyses (Humphries and Reynolds, 2009; Schiller et al., 2011; Horton et al., 2015). Based on function, composition and appearance cell-matrix adhesions can be classified (Sun et al., 2016). Focal adhesions are mainly located near the cell periphery since they establish “early” cell-matrix contacts. They are complex assemblies of many different proteins including the characteristic adapters vinculin, talin, paxillin and the tyrosine-phosphorylated proteins p130Cas, FAK and Src (Zamir and Geiger, 2001). In contrast, the fibrillar adhesions of fibroblasts (previously termed ECM contacts) are found not just at the cell periphery but throughout the entire cell (Geiger and Yamada, 2011; Sun et al., 2016). These fibrillar adhesions are elongated structures that are associated with mature ECM fibrils and are considered to be “late” cell-matrix adhesions. Key components of fibrillar adhesions are extracellular fibronectin, the fibronectin receptor α5β1 integrin and members of the tensin protein family (Zamir et al., 1999; Zamir et al., 2000; Geiger et al., 2001).

Vertebrate tensins represent a family of large cytoskeletal proteins (∼180 kDa) encoded by four genes (Tns1 to Tns4) (Lo, 2004). Tensins 1-3 are focal contact proteins that via their integrin-binding PTB domain at the C-terminus link the cytoplasmic tails of β-integrins to the actin cytoskeleton, bound by their N-terminal actin binding domain (Calderwood et al., 2003). In doing so, tensin 1-3 maintain the tension between in-and outside of a cell. A fourth gene, Cten (=Tns4), encodes a much smaller protein lacking the actin-binding domain of the larger tensin isoforms (Lo, 2004; Haynie, 2014). It has previously been shown that Tns3 is deregulated in cancer and has implications in cell migration, invasion and tumorigenesis, but its role remains controversial (Cui et al., 2004; Katz et al., 2007; Martuszewska et al., 2009; Qian et al., 2009; Cao et al., 2012; Shinchi et al., 2015).

Gaining deeper insights into the plasticity of cell matrix adhesion is of utmost importance to better understand many biological processes including metastasis. In this regard, we desired a cell culture model system reproducing an adhesive and a non-adhesive phenotype of breast cancer cells of the same genetic background. Therefore, we generated a non-adhesively growing (suspension) subclone of the MDA-MB-468 breast cancer cells. Here we report on tensin3 (Tns3) being a major player and target during adhesion plasticity. Identified by transcriptome analysis, Tns3 is dramatically downregulated by loss of cell adhesion. Vice versa knockdown of Tns3 impairs cell adhesion and migration by affecting focal contact composition and dynamics. Moreover, the ectopic expression of Tns3 after loss of cell adhesion (and loss of endogenous Tns3 expression) is able to partially restore cell matrix adhesion and migration. Our results show that the expression as well as the function of Tns3 are tightly intertwined with the cell matrix adhesion in MDA-MB-468 breast cancer cells.

## Results

### Generation and characterization of MDA-MB-468 suspension cells

To gain insights into the plasticity of cell-matrix adhesion and its determinants a culture model consisting of cells which only differ in their mode of adhesion would be a powerful tool. Towards this aim, we have established non-adhesively growing subpopulations of human breast cancer MDA-MB-468 cells. These MDA-MB-468 suspension subclones were selected by repeatedly collecting floating, non-adherent cells from standard MDA-MB-468 cultures and re-cultivating them again on tissue culture dishes. In doing so, a stable suspension subline was established after 12-15 passages. We termed this isogenic suspension subline 468^susp^ while we refer to the adherent parental cell line as 468^par^. The morphology of these two MDA-MB-468 subpopulations is different as revealed by phase contrast microscopy and F-actin staining (Fig. 1A). While 468^par^ cells grow adherently and spread out on tissue culture dishes, 468^susp^ cells grow in suspension and tend to form grape-like cell clusters. Moreover, this adhesion-independent 468^susp^ cell line does not change the suspension growth mode even when provided with adhesive substrates like type I collagen, fibronectin or matrigel (Fig. 1B, and data not shown). Of note, the rate of proliferation and cell death (apoptosis) was found to be fairly similar between both subpopulations, suggesting independence from anoikis induction (Fig. 1C, D).

**Figure 1.**
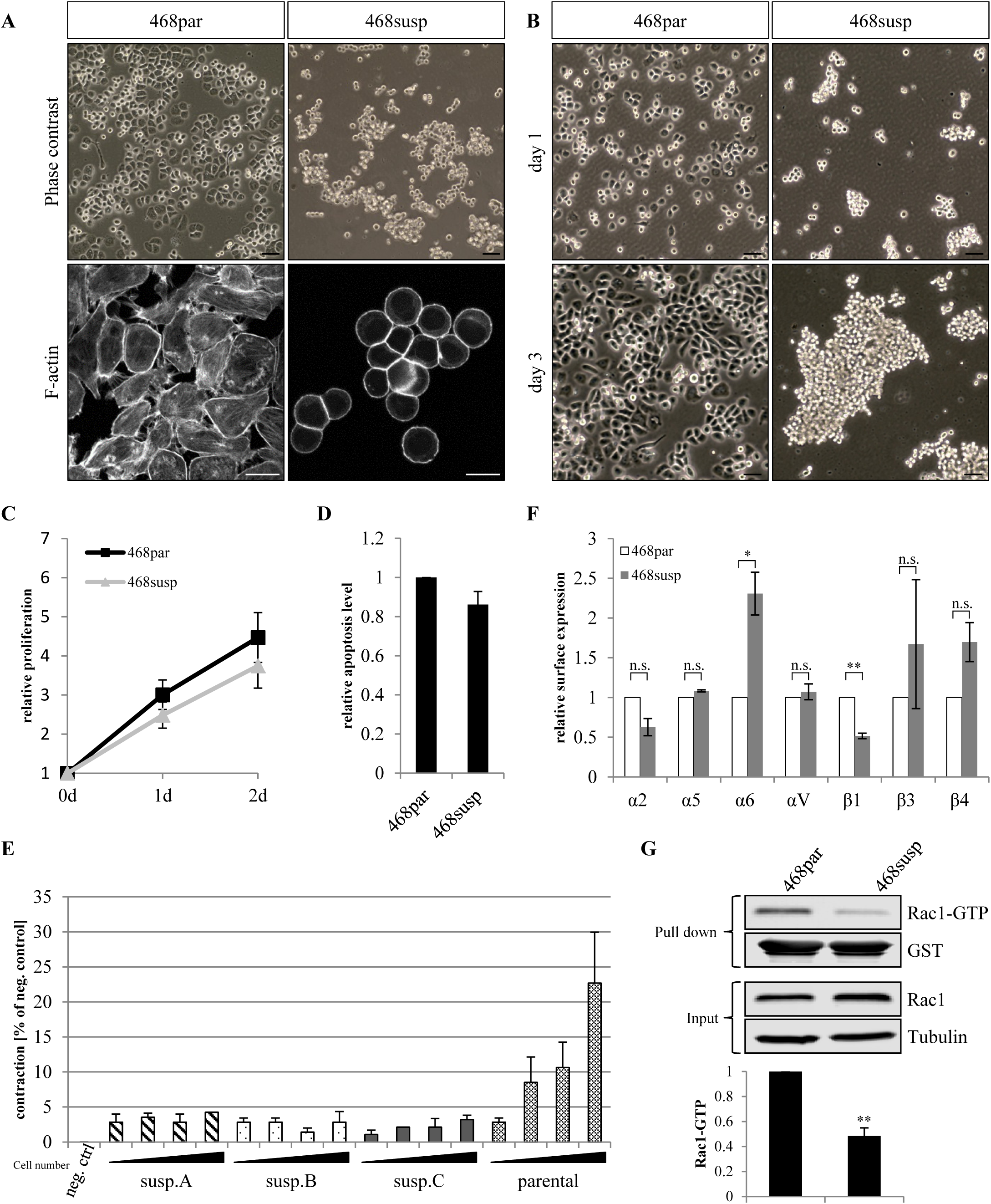
Characterization of MDA-MB-468 subclones selected for non-adhesive growth. **(A)** Representative phase contrast images and F-actin cytoskeleton staining (phalloidin) of parental and suspension MDA-MB-468 breast cancer cells. Scale bars are 50 μm (phase contrast) and 10 μm (F-actin staining). **(B)** Phase contrast images of indicated cells cultured on collagen-coated substrates for indicated time periods. Scale bars, 50 μm. **(C)** Proliferation rate measured in 468^par^ and 468^susp^ cells using a colorimetric assay (n=3). **(D)** Apoptosis rate in 468^par^ and 468^susp^ cells by measuring caspase-3 activity (n=3). **(E)** Collagen contraction assay. Increasing cell numbers of the indicated cell lines were mixed with collagen in medium containing EGF and allowed to solidify in 96 well plates. The area of the collagen lattices was measured after 4 days. Shown is the shrinkage of the collagen gel diameter in % of cell-free negative control (n=3). **(F)** Flow cytometry analysis of integrin surface levels (n=4). **(G)** GST pulldown of Rac1-GTP. Precipitated proteins (GST pulldown) and total lysates (input) were immunoblotted with the indicated antibodies. Quantification of bound active Rac1 normalized to input Rac1 (n=5). All error bars represent ± SEM. P-values were calculated using an unpaired one-sample student’s t-test (*p≤0.05, ** p≤0.01).

Since 468^susp^ cells did not form cell-matrix contacts on flat 2D substrates we examined whether these cells also display impaired adhesion and force transmission within 3D environments. To address this point, we performed a collagen contraction assay. Three independently isolated MDA-MB-468 subclones were encapsulated in 3D collagen hydrogels and the cell-mediated contraction was evaluated after four days. 468^susp^ cells lacked the ability to contract collagen gels independently of the cell concentration (Fig. 1E). In contrast, 468^par^ cells contracted the collagen hydrogels to more than 20% as a function of cell concentration, similar to other adhesively growing cell lines (data not shown). These results show impaired force transmission between the ECM and the actin cytoskeleton in 468^susp^ and indicate defective cell-matrix interactions also in a 3D environment.

The main cell-matrix receptors are integrin heterodimers exposed on the cell surface. Hence, we analyzed by flow cytometry whether the surface expression of different integrin subunits in 468^par^ and 468^susp^ cells is altered. Only integrin α6, β4 and β1 displayed a significant change in surface expression (Fig. 1F). While integrin α6 and β4 was increased, the expression of integrin β1 was decreased in 468^susp^ cells compared to 468^par^ cells. Most integrins, however, were either not expressed (α1, α4, β2; data not shown) or showed no significant difference (α2, α5, αV, β2). Hence, our data indicate that 468^susp^ cells are not defective for integrin presentation but slightly change the subclasses on their cell surface.

A key coordinator in cell adhesion is the small GTPase Rac. It reorganizes the actin cytoskeleton and therefore controls cell behavior including adhesion, spreading and migration. To determine potential differences between parental and suspension cells, the amount and activity of Rac1 protein was determined. Rac1 was significantly less active in 468^susp^ cells compared to the 468^par^ cells as revealed by a PAK-CRIB pull down assay (Fig. 1G). However, immunoblotting of total lysates showed that Rac1 whole protein amounts were comparable.

### Loss of tensin3 in MDA-MB-468 upon loss of adhesion

To investigate the underlying differences between 468^par^ and 468^susp^ cells without bias a genome-wide transcriptome profiling was performed. For this purpose, RNA of 468^par^ cells (3 independent RNA preparations) and of 468^susp^ cells (RNA preparations of 3 independently isolated subpopulations; 468^susp^ A, 468^susp^ B, 468^susp^ C) was used for affymetrix microarray analysis. Using a false discovery rate of less than 5%, we identified more than 300 genes which were significantly up-or downregulated at least twofold in the parental versus suspension cell lines (Suppl. Table 1). Of all genes downregulated in the suspension cells, tensin3 (Tns3) showed the strongest effect, with an average reduction of signal intensity of more than 9-fold in the microarray analysis (Fig. 2A). Tensin3 is an adapter protein that connects the cytoplasmic tail of integrins to the actin cytoskeleton (Calderwood et al., 2003; Lo, 2004). To this end, we thus decided to further analyse the role of Tns3 in adhesion plasticity.

**Figure 2.**
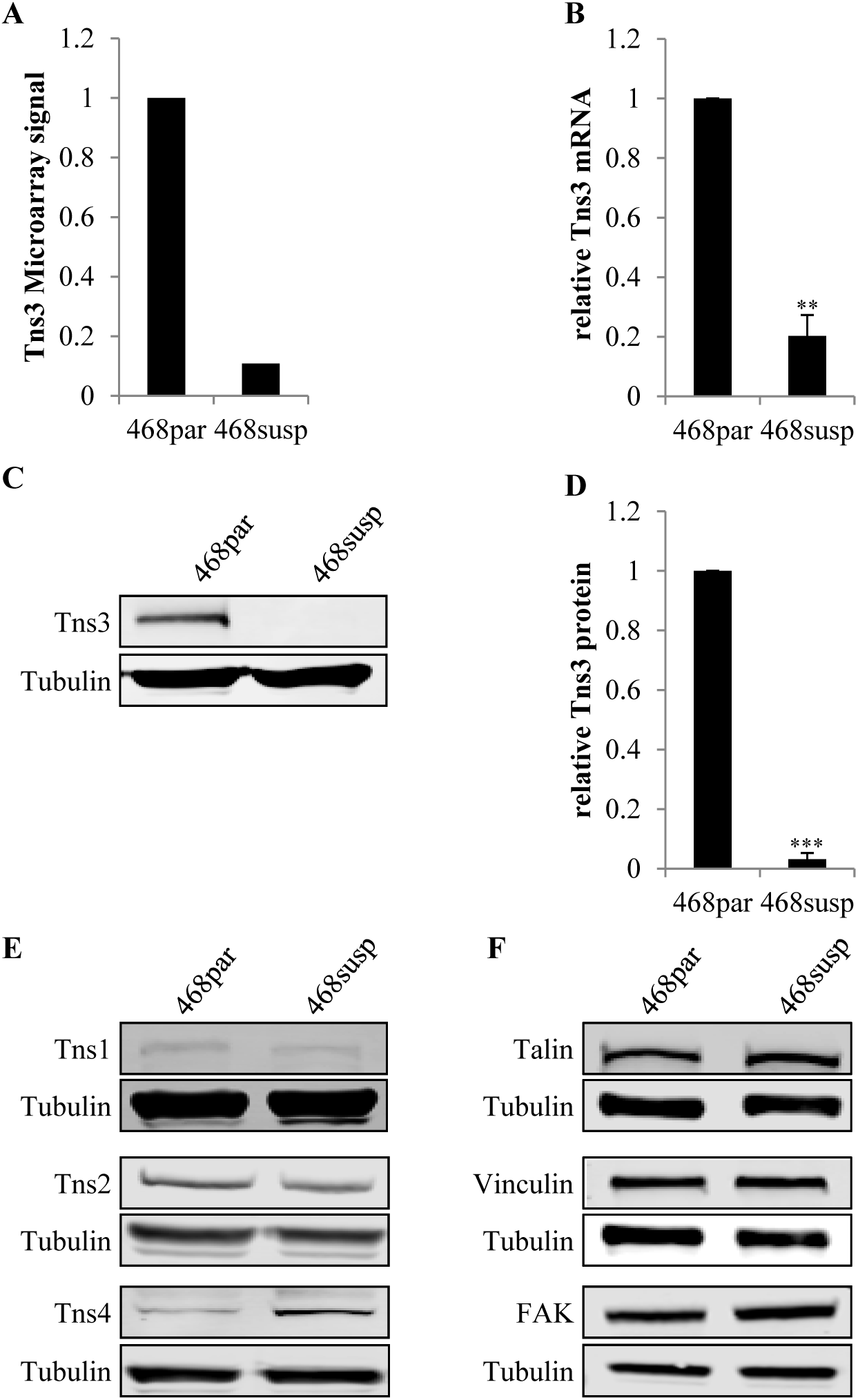
Tns3 expression is decreased in stable suspension subclones. **(A)** Relative microarray signal intensity for Tns3 in MDA-MB-468^par^ and −468^susp^ cells. Shown is the average of 3 independent experiments. **(B)** Tns3 mRNA transcript levels were determined by qRT-PCR normalized to ALAS1 and HPRT1 in 468^par^ and 468^susp^ cells. **(C)** Western blot of Tns3 in total cell lysates from indicated cells. Tubulin was used as loading control. **(D)** Quantification of the Tns3 protein levels normalized to tubulin. **(E)** Representative western blot of Tns1, Tns2 and Tns4 in total cell lysates from indicated cells. Tubulin was used as a loading control. **(F)** Representative western blot of talin, vinculin, and FAK in total cell lysates from indicated cells. Tubulin was used as a loading control. All error bars represent ± SEM, n≥3. All p-values were calculated using an unpaired one-sample student’s t-test (** p≤0.01, *** p≤0.001).

Differential expression of Tns3 in 468^susp^ cells compared to 468^par^ cells was validated by quantitative RT-PCR which showed a 6-fold decrease of Tns3 mRNA level in 468^susp^ cells (Fig. 2B). However, the tensin3 protein was almost undetectable in 468^susp^ cells by immunoblotting (Fig. 2C). Quantification revealed a 30-fold downregulation in the suspension cells (Fig. 2D). In comparison, the transcript and protein level of other tensin family members were also analyzed by qRT-PCR and western blot analysis. The transcript level of Tns2 was downregulated, whereas Tns1 and Tns4 mRNA amounts were increased in 468^susp^ cells (Suppl. Fig. S1A). However, the protein level of Tns1 and Tns2 were unaffected, whereas Tns4 protein was increased in 468^susp^ cells (Fig. 2E). We further investigated if the expression levels of other major focal adhesion adapter proteins are also affected in 468^susp^ cells. Interestingly, we observed no decrease on the protein or transcript level of other focal adhesion proteins like vinculin, FAK or talin (Fig. 2F, Suppl. Fig. S1B, C). However, immunofluorescence staining revealed that these proteins do not localize to focal adhesion like structures in 468^susp^ cells, consistent with their lack of focal adhesions (data not shown).

To elucidate whether loss of cell-matrix adhesion is sufficient to induce downregulation of Tns3 in 468^par^ cells, these cells were cultivated on non-adhesive polyHEMA-coated substrates for up to 72 hours (Fig. 3). Unlike the stably selected 468^susp^, these cells however restored matrix adhesion after non-adhesive growth when transferred back to tissue culture plastic (Fig. 3A). By this experiment we could show that transient loss of adhesion resulted in a significant decrease of Tns3 in MDA-MB-468 (Fig. 3B, C). Within the first 24 hours Tns3 transcript and protein levels dropped to around 60% and reached less than 40% of their original expression after 72 hours. Importantly, readhesion restored Tns3 in MDA-MB-468 on both protein and mRNA level to almost 100% within 24 hours. Similar results were obtained with MCF7 breast cancer cells (Suppl. Fig. S2). However, in MCF7 cells the dynamics are slightly different as Tns3 levels decreased stronger and are just partially restored within 48 hours of readhesion. In contrast to tensin3 the protein levels of talin and vinculin were not significantly altered by cultivating MDA-MB-468^par^ or MCF7 cells on non-adhesive polyHEMA substrate (Fig. 3D, E and Suppl. Fig. S2D, E).

**Figure 3.**
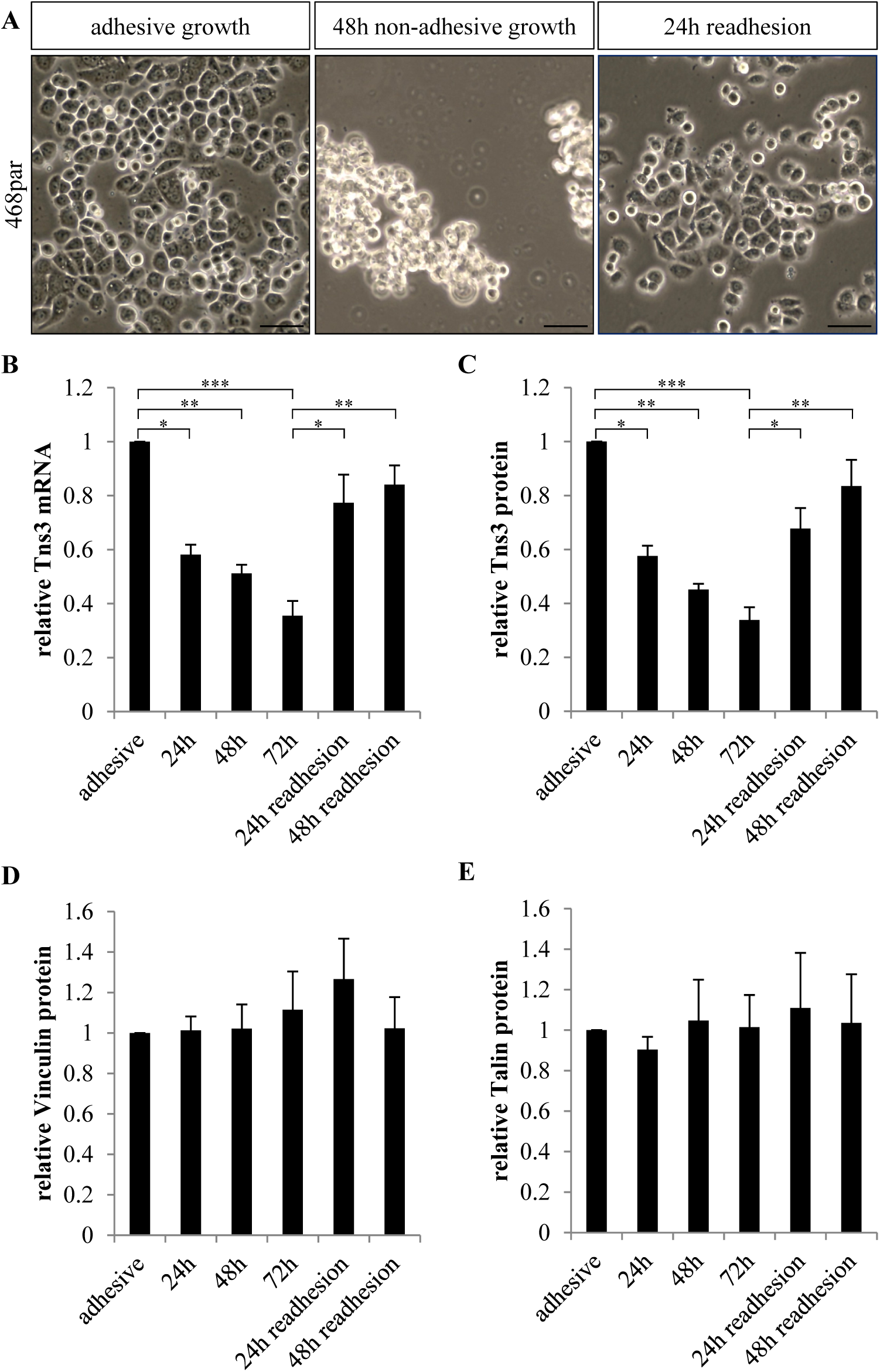
Adhesion-dependent expression of Tns3 in MDA-MB-468 cells. **(A)** 468^par^ cells were non-adhesively cultivated on polyHEMA coated dishes for up to 72 h. Afterwards cells were allowed to re-adhere to normal culture dishes. Phase contrast micrographs at the times indicated. Scale bars, 50 μm. **(B)** Tns3 mRNA transcript levels were determined by qRT-PCR and normalized to ALAS1 and HPRT1 expression. **(C)** Quantification of Tns3 protein levels in total cell lysates from indicated conditions normalized to tubulin. **(D)** Quantification of vinculin protein levels in total cell lysates from indicated conditions normalized to tubulin. **(E)** Quantification of talin protein levels in total cell lysates from indicated cells normalized to tubulin. All error bars represent ± SEM, n=3. All p-values were calculated by Tukey’s post-hoc test using one-way ANOVA (* p≤0.05, ** p≤0.01, *** p≤0.001).

### Tensin3 knockdown influences adhesion and migration capacity of 468^par^ cells

To address whether loss of Tns3 alone interferes with cell-matrix adhesion of MDA-MB-468 cells, stable knockdown of Tns3 in 468^par^ cells was performed. Therefore, 468^par^ cells were lentivirally transduced with three independent constructs encoding Tns3 specific shRNAs or a control shRNA. Expression of Tns3 specific shRNAs resulted in up to 80% decreased Tns3 levels (Fig. 4A, B). To gain further insights into the defects of 468^par^ cells lacking Tns3 we performed cell adhesion assays. In these assays equal numbers of knockdown cells were plated on type I collagen or fibronectin coated wells. After allowing adhesion and spreading for 30 and 60 min, a profound decrease in the number of attached cells was observed for Tns3-depleted cells (Fig. 4C). This decrease was observed by all applied Tns3 shRNAs and was evident on both substrates. A similar albeit less pronounced decrease of cell adhesion was also seen in HCT8 colon cancer cells upon stable Tns3 knockdown (Suppl. Fig. S3). The MDA-MB-468 knockdown cells which remained attached lacked Tns3-positive cell-matrix adhesions and covered a smaller area on collagen than the control cells, suggestive of an impaired spreading behavior (Fig. 4D, E). The reduced cell size could be verified by transient knockdown using specific Tns3 siRNAs (Suppl. Fig. S4). In addition, we performed a transwell migration assay to examine the migratory capacity. Even without chemotactic stimulation, the undirected migration of Tns3 depleted cells was slightly but significantly decreased (Fig. 4F). Using EGF as a chemotactic stimulus, we further observed that the migratory activity of Tns3 knockdown cells was strongly decreased compared to control cells. Hence, knockdown of Tns3 impaired migration of MDA-MB-468 cells in both unstimulated and EGF-stimulated conditions.

**Figure 4.**
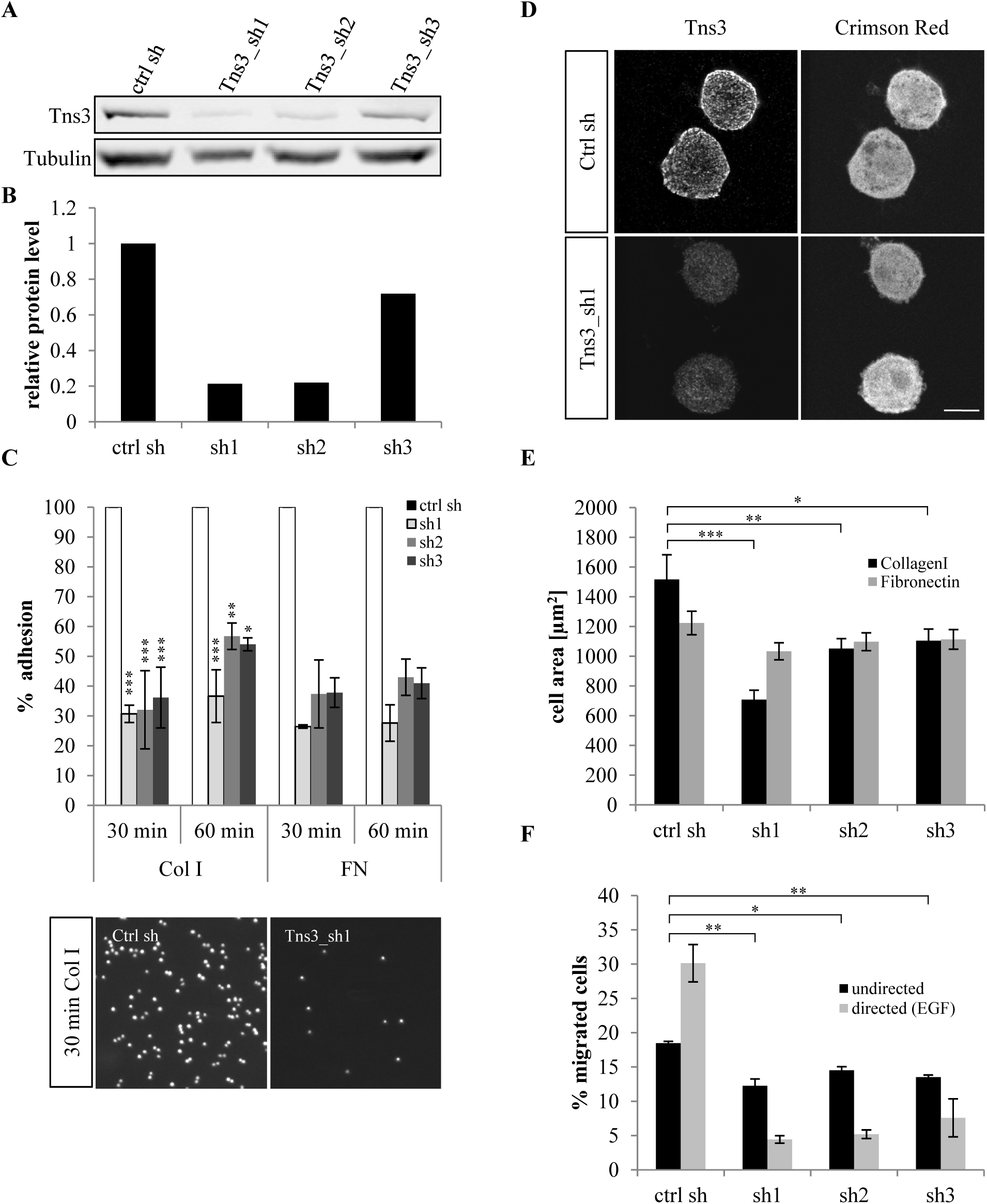
Adhesion and migration of MDA-MB-468 cells upon stable Tns3 knockdown. Three different Tns3 targeting shRNA constructs (Tns3_sh1, _sh2, _sh3) were lentivirally transduced and cells were selected with puromycin (2 μg/ml). As control a non-targeting sequence (ctrl sh) was used. **(A)** Representative western blot showing levels of total Tns3 and tubulin in stable cell lines. **(B)** Quantification of Tns3 protein levels in total cell lysates from indicated cells normalized to tubulin. **(C)** Adhesion of stable cell lines to collagen (Col I)- or fibronectin (FN)-coated surfaces for the indicated times. Shown is the percentage of adhered cells normalized to the control (Col: n=3, FN: n=2). Representative DAPI images of cells plated on type I collagen for 30 min before fixation are shown below. **(D)** Representative images of cells cultivated overnight on collagen, followed by immunostaining of Tns3. Stable transduction is indicated by Crimson red expression. Scale bars, 10 μm. **(E)** Cell area determination by automated tracking of 50-100 cells per condition. Cells were seeded on coated 8-well chamber slides for 30 min before spreading of individual living cells was monitored over 15 h. **(F)** Transwell cell migration for 22 h with or without EGF as a chemoattractant, normalized to the seeding controls. The error bars represent ± SEM, n=3. All p-values were calculated by Tukey’s post-hoc test using one-way ANOVA (*p≤0.05, ** p≤0.01, *** p≤0.001).

To examine the underlying structural changes of cell-matrix contacts, we analyzed the immunofluorescence pattern of the focal adhesion proteins vinculin, talin and paxillin in cells which adhered despite Tns3 knockdown. For this purpose, Tns3-depleted and control cells were allowed to adhere and spread out for 15 min, 30 min and 60 min on collagen (Fig. 5) or fibronectin coated coverslips (data not shown). In cells transfected with a control shRNA, the proteins vinculin and paxillin were detected in the typical dot-like focal adhesions at the tip of F-actin fibers (Fig. 5A). Methanol/acetone fixation also revealed a similar localisation for talin (Fig. 5B). All three focal adhesion proteins were located at the moving front of the spreading control cells. In contrast, the Tns3 depleted cells displayed an altered localization and appearance of focal adhesion proteins, and diminished spreading. Vinculin, paxillin and also talin appeared to mainly accumulate at peripheral positions in thick focal adhesion plaques in Tns3 knockdown cells (Fig. 5, right panels). Colocalising with the deformed adhesion sites, Tns3 depleted cells showed strong cortical F-actin staining around the cell periphery. In those Tns3 depleted cells that still adhered to the matrix the further processing of the initial focal adhesions into more mature structures seems impaired.

**Figure 5.**
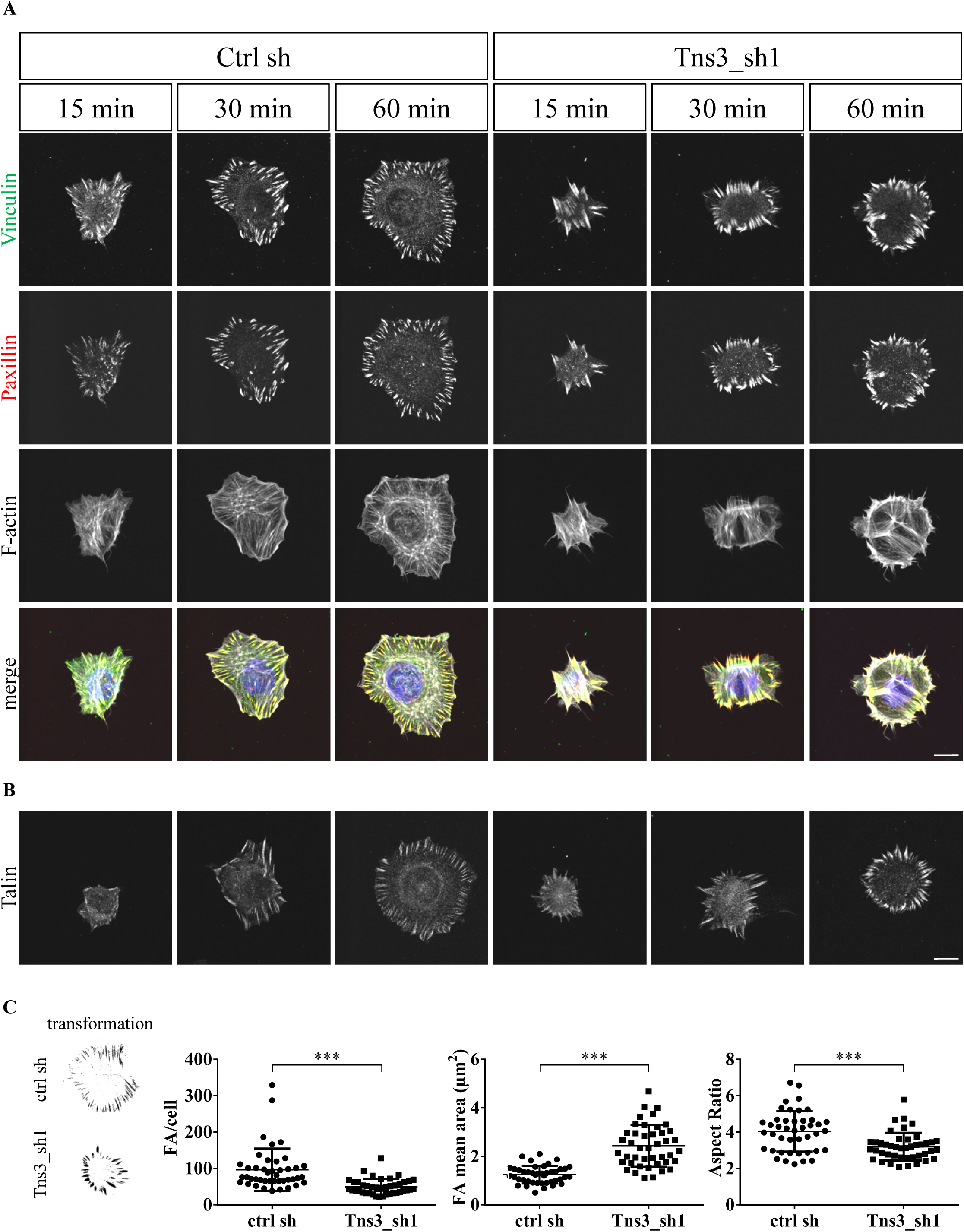
Tns3 knockdown results in altered focal adhesion formation. Stable Tns3 knockdown cells (Tns3_sh1) and ctrl shRNA cells were plated on collagen coated glass slides, stained for the indicated proteins and analyzed by fluorescence confocal microscopy. Representative images are shown. Scale bars, 10 μm. **(A)** Co-immunostaining of vinculin (green), paxillin (red) and F-actin (SiR-Actin, depicted in white). Cells were fixed by formaldehyde 15, 30 and 60 min after seeding. Merge images show false color overlay (blue: DNA). **(B)** Talin immunostaining following MeOH/acetone fixation. **(C)** Number, size and aspect ratio of paxillin-positive focal adhesions from 43 Tns3 knockdown and control cells after 60 min of adhesion. Calculations were done after binarized image transformation (examples in left panels) and automated measurement. Scatter plots show FA number, mean area and mean aspect ratio per cell, with average ± SD indicated. Statistical significance according to unpaired two-sample student’s t-test (*** p≤0.001).

Automated measurements of paxillin-positive structures showed a reduction of focal adhesions per cell to around 50% upon Tns3 knockdown (Fig. 5C). Whilst the mean area covered by individual focal adhesions significantly increased, the aspect ratio decreased, indicative of abnormal bundling or thickening of the adhesion sites (Fig. 5C). Together, these results suggest that reduction of Tns3 affects the dynamic of focal adhesion remodelling during spreading. This defect in focal adhesion maturation may be responsible for rounding and the reduced size of Tns3 deficient cells.

### Tensin3 rescue restored adherent growth of 468^susp^ cells

We have shown that Tns3 is directly regulated by the status of adhesion and is lost in non-adherent 468^susp^ cells. Next, we asked if exogenous expression of full-length Tns3 is able to restore adherent growth of the 468^susp^ cells. To address this question, suspension cells were transfected with plasmids containing either a GFP-Tns3 or a GFP sequence followed by Geneticin (G418) selection. Immunoblot analysis revealed that selected cells showed an approximately threefold overexpression of Tns3 compared to the parental cells (Fig. 6A). During the selection procedure both flotile and adherent cells were cultivated. After 3 weeks of selection the number of adherently growing cells was determined. Importantly, we found about 20 fold more adherently growing cells expressing GFP-Tns3 than expressing GFP alone (Fig. 6B). Despite potential overexpression artefacts, this result suggested that re-expression of Tns3 in 468^susp^ is sufficient to revert the non-adherent phenotype of these cells. Moreover, the localization of Tns3-GFP into focal contacts was indistinguishable from the 468^par^ cells as shown by confocal microscopy (Fig. 6D). Furthermore, 468^susp^ cells stably overexpressing Tns3-GFP displayed a similar F-actin cytoskeleton architecture compared to the 468^par^ cells and were able to restore the formation of F-actin fibres as well as the localization of vinculin to focal adhesion like structures (Fig. 6D). In contrast, 468^susp^ cells completely failed to attach and hence any focal contacts were absent (data not shown).

**Figure 6.**
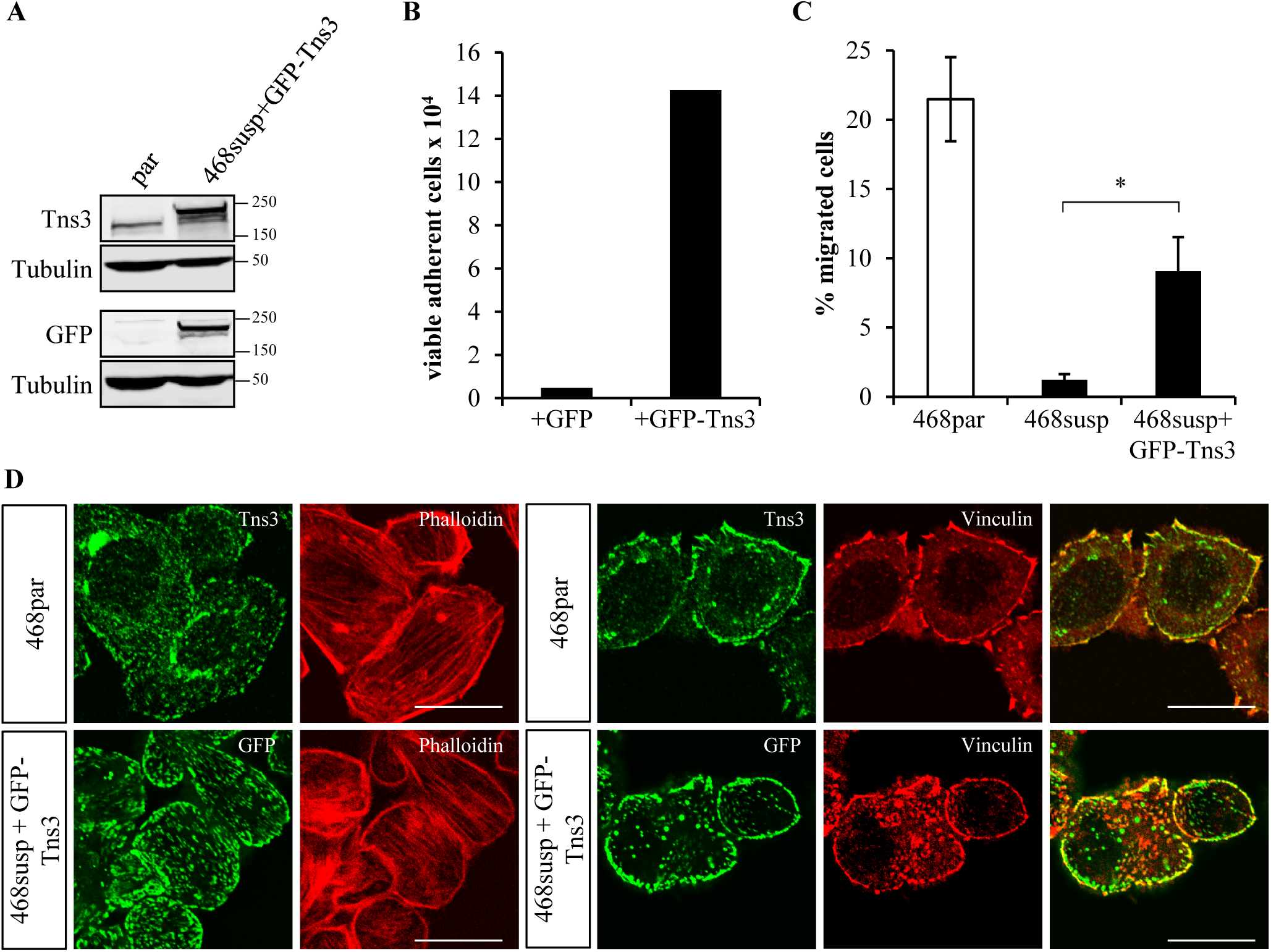
Tns3 overexpression restored adherent growth of 468^susp^ cells. MDA-MB-468^susp^ cells were stably transfected with GFP-tagged Tns3 or with the pEGFPc1 vector as a control. During the selection procedure both flotile and adherent cells were cultivated. **(A)** Representative immunoblots of total cell lysates using Tns3-and GFP-specific antibodies. Tubulin was used as a loading control. **(B)** Number of adherently growing cells after 3 weeks of selection. **(C)** Transwell migration of 468^par^, 468^susp^, and 468^susp^ cells expressing GFP-Tns3, measured after 22 h with EGF as a chemoattractant and normalized to the seeding controls. All error bars represent ± SEM, n=4. * p≤0.05 according to unpaired two-sample student’s t-test **(D)** 468^par^ and 468^susp^ cells expressing GFP-Tns3 were stained for Tns3 (green) together with F-actin (phalloidin, red) or vinculin (red, right panels). Merged images show endogenous Tns3 or GFP-Tns3 (green) and vinculin (red). Scale bars, 10 μm.

Cell-matrix adhesion is an important feature of cell migration and motility. Therefore, we next analyzed whether the Tns3-GFP mediated re-adhesion alters the migratory activity of 468^susp^ cells by applying a transwell assay experiment. The cell migration of 468^susp^ cells stably expressing Tns3-GFP was significantly increased compared to the 468^susp^ control cells (Fig. 6C). Compared to 468^par^ cells, however, cell migration was only partially restored by Tns3-GFP, possibly explainable by either overexpression of Tns3-GFP or additional alterations in the selected suspension cells. Nevertheless, our results showed that Tns3-GFP expression restores cell adhesion of 468^susp^ cells and enhance the migratory activity of these cells.

## Discussion

### Generation of MDA-MB-468 suspension cells

In this study, we report on the regulation and role of the cell-matrix contact protein Tns3 in breast cancer cells during adhesion plasticity. We show that Tns3 expression is directly correlated with the status of cell adhesion and is lost following loss of cell-matrix adhesion in MDA-MB-468^susp^ cells, an enriched and stable subpopulation of MDA-MB-468 breast cancer cells growing in suspension.

Culturing cells in or on non-adhesive substrates can induce adhesion-independent growth of typical adhesively growing cells such as epithelial cells. In doing so cells are forced to grow without substrate attachment and cell-matrix contacts which results into a constrained suspension phenotype. However, in our study we aimed at establishing a method that considers the inherent status of cell-matrix adhesion and still reflects on adhesion plasticity, i.e. shows adherent and non-adherent growth. Towards this end, we have successfully generated non-adherently growing suspension sublines of MDA-MB-468 breast cancer cells which don’t display obvious signs of genetic drift, as confirmed by identical STR analysis (data not shown). These suspension sublines were acquired by enriching non-adherent single cells that spontaneously dissociated from MDA-MB-468 monolayer cultures. In a similar approach Tang et al. generated an in vitro metastasis model consisting of metastasis-like dissociated round cells (R-cells) and adherent epithelial cells (E-cells) from HCT8 colon adenocarcinoma cells (Tang et al., 2010; Tang et al., 2014). In HCT8 colon cancer, R-cells occurred at very low frequency but the R-E ratio could be increased by appropriate physical properties of the culture substrate. In our study we were able to achieve stable suspension MDA-MB-468 sublines after repeatedly separating and enriching flotile cells. It is remarkable that this flotile cell population is neglected in most studies with MDA-MB-468 since non-adherent cells normally are discarded by medium aspiration. Presumably, also other cancer cell lines contain flotile subpopulations that are not taken into account by many experimental approaches. In contrast, we place the suspension phenotype of MDA-MB-468 cells into the spotlight and utilize it to investigate adhesion plasticity in human breast cancer cells.

After losing cell-matrix adhesion or sensing inappropriate cell-matrix contacts epithelial cells normally undergo apoptotic cell death; a process known as anoikis (Frisch and Francis, 1994; Frisch and Screaton, 2001; Gilmore, 2005). In tissue homeostasis anoikis can be seen as a self-defense strategy that prevents cells loosing ECM contacts from dysplasia formation and metastatic dissemination (Guadamillas et al., 2011). However, many cancerous cells are able to avoid anoikis and have developed anoikis-resistance (Frisch and Screaton, 2001; Simpson et al., 2008; Taddei et al., 2012; Paoli et al., 2013; Buchheit et al., 2015). In this regard, the generation of non-adherent 468^susp^ was achievable because MDA-MB-468 cells are cancer cells whose loss of SMAD4 and subsequent caspase signaling was proposed to render them anoikis-resistant (Ramachandra et al., 2002). Furthermore, we observed in both 468^susp^ and polyHEMA-induced suspension of 468^par^ that cells form prominent cell-cell contacts and grow in grape-like structures. Tight cell-cell contact formation is a typical phenomenon of epithelial cells in suspension and has been discussed as compensatory mechanisms to prevent apoptosis after loss of cell-ECM adhesion (Kantak and Kramer, 1998; Zhan et al., 2004; Hofmann et al., 2007). Notably, although the protein levels of cell-cell contact protein E-cadherin are fairly similar E-cadherin localizes more prominently to cell-cell contacts in 468^susp^ than in 468^par^ (Suppl. Fig. S5C, D). Moreover, an adaptation/compensation of 468^susp^ towards more robust anoikis-resistance is indicated also by the increased expression of the cell-cell contact protein CEACAM6 (Suppl. Fig. S5A, B) which has been shown to increase anoikis-resistance in carcinoma cells (Ordonez et al., 2000; Duxbury et al., 2004).

### Tensin3 is lost in MDA-MB-468 suspension cells and directly regulated by cell-matrix adhesion

To examine the impact of adhesion plasticity on gene regulation in breast cancer cells we tested the differential gene expression between 468^par^ and 468^susp^ by microarrays. We found that the cytoskeletal adapter protein tensin3 (Tns3) is strongly reduced in suspension cells on both mRNA and protein level. Loss of Tns3 mRNA and protein is directly caused by loss of cell-matrix adhesion but restored by gain of cell-matrix adhesion as corroborated by short-term cultures of 468^par^ on non-adhesive polyHEMA substrate. Furthermore, the adhesion-mediated regulation of Tns3 is not restricted to MDA-MB-468 cells since other breast cancer cells such as MCF7 lose Tns3 expression after loss of cell-matrix adhesion as well (Suppl. Fig S3). Although it seems reasonable that non-adherent cells downregulate focal adhesion proteins our data clearly show that other major cell-matrix adapter proteins such as vinculin and talin are not affected by loss of cell-matrix adhesion. This result might be explained by the proposed role of Tns3 in “late” cell-matrix contacts while vinculin and talin are important in “early” and “intermediate” focal adhesions (Zamir et al., 1999; Katz et al., 2000; Zamir et al., 2000; Zaidel-Bar et al., 2003; Zaidel-Bar et al., 2004). In this regard, it is conceivable that cells retain the expression of “early” adhesion proteins while they shut down the expression of “late” adhesion proteins missing the right chemical or physical signal from “early” adhesion. However, our results show less dramatic effects for Tns2 and Tns1. Hence, the adhesion mediated regulation of Tns3 might be an isoform-specific effect.

We have demonstrated that cell-matrix adhesion regulates Tns3 expression. Conversely, we show that Tns3 itself influences the adhesion properties of MDA-MB-468. By knockdown experiments we show that matrix adhesion including cell spreading dynamics and focal adhesion composition are impaired in 468^par^ after stable shRNA-mediated Tns3 knockdown. In Tns3-deficient cells which still adhere, the proper adhesion maturation from “early” into “late” adhesions seems impeded. This finding is in accordance with work from Pankov et al. where the authors showed by disrupting the tensin function in human fibroblasts that “late” fibrillar adhesions disappeared but the “early” focal adhesions stayed intact indicating that tensin is necessary for adhesion maturation (Pankov et al., 2000). Furthermore, we found that knockdown of Tns3 decreases the migratory activity in MDA-MB-468 which could be related with the impeded adhesion maturation. This result coincides with work from Qian et al. and Shinchi et al. that reported reduced cell migration in four carcinoma cell lines including MDA-MB-231 and MDA-MB-468 after Tns3 knockdown (Qian et al., 2009; Shinchi et al., 2015). In contrast, Katz et al. and Cao et al. showed that Tns3 knockdown in non-tumorigenic mammary MCF 10A cells resulted into an increased migration behavior (Katz et al., 2007; Cao et al., 2012). In these studies, EGF treatment was proven to strongly increase the migration of MCF 10A (Katz et al., 2007; Cao et al., 2012). Similarly, EGF induced cell migration in MDA-MB-468 cells in our experiments. However, it was unexpected that EGF intensified the Tns3 knockdown mediated decrease in cell migration in MDA-MB-468. In this context, it is important to note that EGF itself has been shown to decrease Tns3 but increase Tns4 (Cten) levels in MCF 10A and therefore causes a Tns3-to-Tns4 switch in these non-tumorigenic cells (Katz et al., 2007). Remarkably, we found a Tns3-to-Tns4 switch in MDA-MB-468 as well. However, the Tns3-to-Tns4 switch in MDA-MB-468 cells occurred independent of EGF treatment but as a function of cell-matrix adhesion. While loss of cell-matrix adhesion decreased Tns3 levels in 468^susp^ it significantly increased the Tns4 mRNA and protein levels. Interestingly, a 7-fold increase in Tns4 induced by loss of cell-matrix adhesion has been shown as well in the HCT8 colon cancer R-E model from Tang et al. (Tang et al., 2014). We observed that Tns3 knockdown reduced matrix adhesion of HCT8 cells (Suppl. Fig. S3), but it is unknown whether Tns3 is altered in the R-E model. Whether the adhesion-mediated and the EGF-mediated Tns3-to-Tns4 switches share the same molecular downstream mechanisms needs to be addressed in further studies.

### Tensin3 is target and regulator of cell-matrix adhesion

Tns3 directly links β-subunit integrins with the F-actin cytoskeleton (Lo, 2004) and has been obviously named by its key function: maintaining cellular tension. In our study we show that not just Tns3 is responsible for maintaining proper cell-matrix adhesion (and therefore tension) but that vice versa cell-matrix tension is crucial for Tns3 expression. Hence, we propose a Tns3 feedback loop in which proper cell-matrix adhesion and Tns3 expression are tightly intertwined in breast cancer cells (see Fig. 7). Loss of cell-matrix adhesion results into the disappearance of Tns3. In turn, loss of Tns3 impairs cell-matrix adhesion as shown by our Tns3 knockdown experiments and together with other changes maintains loss of cell-matrix adhesion. Once cells have left the Tns3 feedback loop and lost cell-matrix adhesion Tns4 becomes upregulated. Although Tns4 is able to bind to the β-subunit of integrins it lacks the actin-binding domain of the larger tensin isoforms (Lo and Lo, 2002; Lo, 2004) and therefore is unable to mediate cell-matrix adhesion. Hence, it is conceivable that the upregulated Tns4 might contribute to the suppression of Tns3 expression and keeps suspension cells outside the Tns3-feedback loop. In addition, we have shown that cell-matrix adhesion can be partially restored in flotile 468^susp^ cells by GFP-Tns3 overexpression cells. This finding corroborates our conclusion that Tns3 and cell-matrix adhesion are mutually dependent in MDA-MB-468 breast cancer cells. It remains to be investigated whether such a mutual regulation plays a role during cycles of metastatic dissemination through the bloodstream and colonization at distal sites, when cancer cells diminish and regain adhesive properties, respectively. Moreover, although tensins have been originally identified as a protein family with structural functions, Tns3 is also an important docking platform for signaling molecules. In this regard, Tns3 has been shown to bind several pro-oncogenic tyrosine phosphorylated proteins, including p130Cas, Sam68, Src and FAK (Cui et al., 2004; Defilippi et al., 2006; Mitra and Schlaepfer, 2006; Qian et al., 2009). Additionally, Tns3 is also able to bind DLC1 (deleted in liver cancer 1) and Dock5 which are both components of the Rho GTPase pathways (Cao et al., 2012; Touaitahuata et al., 2016; Blangy, 2017). Therefore, it seems likely that the adhesion-mediated Tns3 regulation entails strong consequences on downstream signaling involved in the fate of cancer cells.

**Figure 7.**
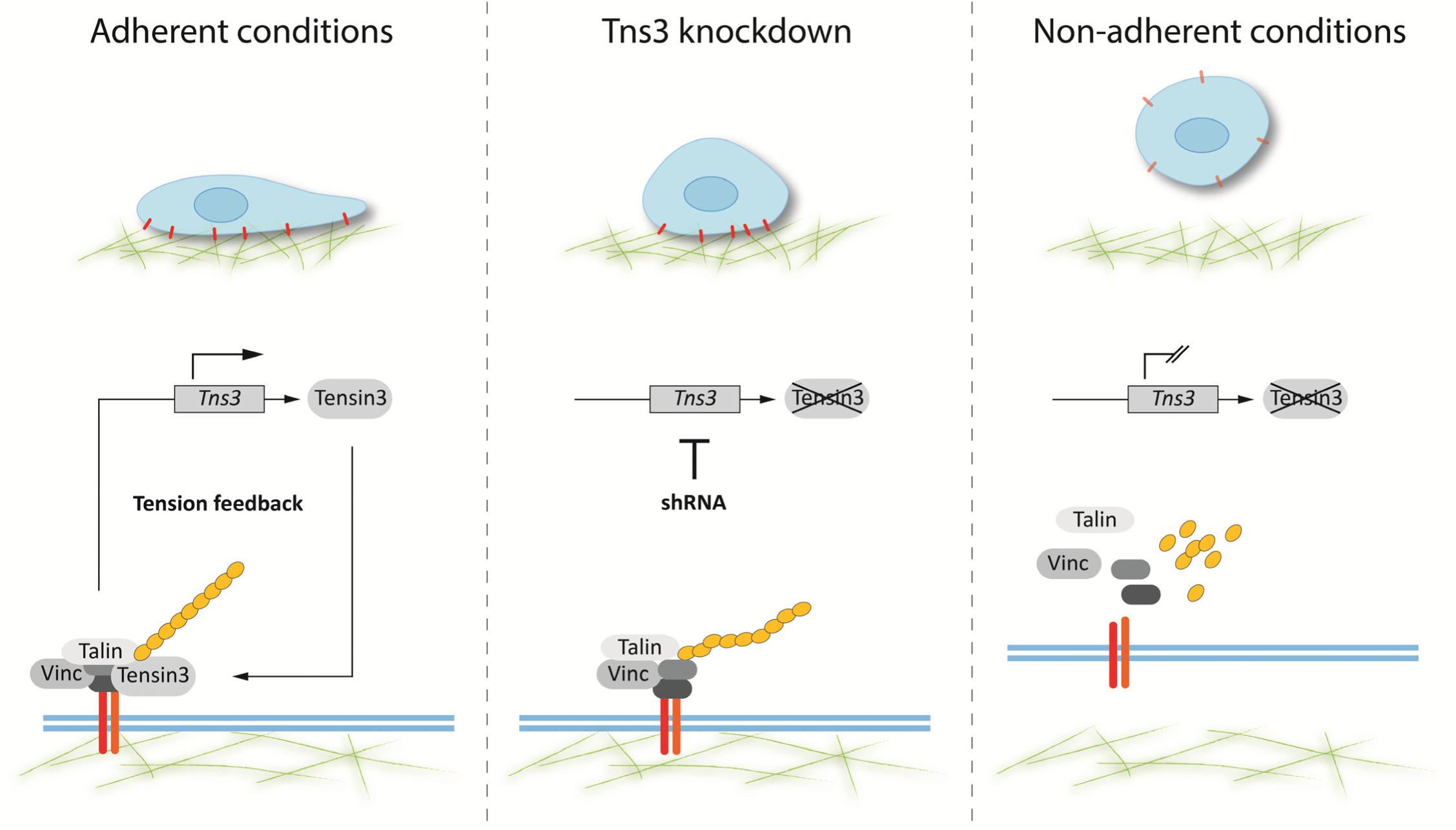
Model of the interplay between Tns3 expression and cell-matrix adhesion. During adherent conditions cells attach via integrin-containing cell-matrix contacts (red) to the substrate (green fibers). This results in a positive feedback signal necessary for Tns3 expression. In turn, Tns3 protein is expressed and contributes to proper cell-matrix contacts by connecting integrins (orange, red) to the actin cytoskeleton (yellow). This feedback loop can be disturbed by either loss of Tns3 expression or loss of cell-matrix adhesion. Loss of Tns3 expression by i.e. shRNA leads to the disappearance of the Tns3 protein. Consequently, cell adhesion, migration and focal adhesion formation are impaired. On the other side, loss of cell-matrix adhesion prevents the required positive feedback from adhesion and results into a shutdown of the Tns3 gene expression and protein production.

## Materials and Methods

### Plasmids and Reagents

Human full length Tns3 was cloned into pEGFPc1 (Clontech, Mountain View, CA) for mammalian expression. ShRNA constructs (Tns3_sh1: 5’-GCATCACCCTGACAGACAATCTTCAAGAGAGATTGTCTGTCAGGGTGATGCTTTTTTACGCGT-3’, Tns3_sh2: 5’-GG ACGCATAGGAGTGGTCATATCATTTCAAGAGAATGATATGACCACTCCTATGCGTCCTTTTTTACGCGT-3’, Tns3_sh3: 5’-GAAAGCTGGAGATTTGGCCAATGAATTCAAGAGATTCATTGGCCAAATCTCCAGCTTTCTTTTTTACGCGT-3’, ctrl_sh 5’-TTGTACTA CACAAAAGTACTGTTCAAGAGACAGTACTTTTGTGTAGTACAATTTTTTACGCGT-3’) were cloned in pLVX-shRNA2 Crimson kindly provided by Stefan Hüttelmaier. This vector is based on pLVX-shRNA2 (Clontech Laboratories, Mountain View, CA). The ZS Green was replaced by E2-Crimson, a far-red fluorescent protein. The siRNAs were purchased as siRNA duplexes from Sigma-Aldrich (Steinheim, Germany). The sequences were: siTns3-1 5’-CCAGGCCCUUGACAGGUUU-3’, siTns3-3 5’-GGACAAUGCCAGCAAAGAU-3’, siTns3-5 5’-GGUUGUAGCUCACCAGUAU-3’ and siTns3-9 5’-GCUCGAAUGAACCAUAUUU-3’. As control *Silencer* Negative Control #1 siRNA (Thermo Fisher Scientific, cat. no. AM4636) was used. For microarray analysis the Human Gene ST 1.0 array, comprising 28.869 probe sets covering the entire transcriptome was used. The labeling, hybridization and quality control was carried out as described (Descot et al., 2009). The data were analyzed in collaboration with Reinhard Hoffmann (TU Munich), using RMA normalisation and the permutation-based SAM algorithm. The q-value is the lowest false discovery rate at which the gene is called significant, applying a threshold of 4.62%. Microarray datasets have been deposited at the Gene Expression Omnibus database (accession number GSE98062).

Primary antibodies used were anti-GFP (Sigma, catalog no. G1544), anti-GST (Sigma, catalog no. G7781), anti-vinculin (Sigma, catalog no.V9131), anti-talin (Sigma, catalog no. T3287), anti-tubulin (Sigma, catalog no. T9026), anti-Tns3 (Millipore, catalog no. ABT29), anti-FAK (Millipore, catalog no. 06-543), anti-Tns1 (abcam, catalog no. ab135882), anti-paxillin (abcam, catalog no. ab32084), anti-Tns2 (Cell Signaling, catalog no. 11990), anti-Tns4 (Abnova, catalog no. H00084951-MO), anti-Rac1 (BD bioscience, catalog no. 610650), anti-E-cadherin (BD Bioscience, catalog no. 610181), anti-integrinß1, ß3, ß4 and α2, α5, α6, αV (for flow cytometry all from Miltenyi Biotec, Bergisch Gladbach, Germany). F-actin was visualized with Atto 546 Phalloidin (Sigma-Aldrich) or SiR-Actin (Cytoskeleton, Denver, CO, USA). Alexa Fluor-conjugated and IRDye^®^ (800CW, 680RD) secondary antibodies were from Thermo Fisher Scientific or Li-Cor (Bad Homburg, Germany), respectively. DNA was stained with DAPI (Sigma-Aldrich).

### Cell Culture and Transfection

The breast cancer cell line MDA-MB-468 was cultured at 37 °C and 5% CO_2_ in RPMI medium supplemented with 10% (v/v) fetal calf serum (FCS) and antibiotic/antimycotic (Thermo Fisher Scientific, Schwerte, Germany). For non-adhesive growth cell culture dishes were coated with 1.2% (w/v) polyHEMA (Poly(2-hydroxyethyl methacrylate, Sigma-Aldrich) dissolved in ethanol. Dishes were air-dried for 12 h in a laminar flow hood under sterile conditions. MCF7 cells were grown in Dulbecco’s Modified Eagle Medium (DMEM) high glucose supplemented with 10% (v/v) fetal calf serum (FCS), 1% sodium pyruvate (v/v) and antibiotic/antimycotic.

Transfections were carried out using Lipofectamine LTX (Thermo Fisher Scientific). 1 μg plasmid DNA was added to 5x10^5^ 468^susp^ cells (12-well plates) according to the manufacturer’s instructions. Cells were selected using 1 mg/ml Geneticin (G418, Thermo Fisher Scientific). Lentiviral transductions were done by adding concentrated virus titer and 10 μg/ml polybrene to 3x10^5^ cells (12-well plates) for 24 h. Cells were selected using 2 μg/ml puromycin (Thermo Fisher Scientific). For virus production, HEK293T cells were cotransfected by CaPO_4_ precipitation with pMD2.G (Addgene, no. 12259), psPAX2 (Addgene, no. 12260) both kindly provided by Stefan Hüttelmaier and pLVX-shRNA2 crimson containing either ctrl shRNA or Tns3 targeting sequence. Lentiviruses were purified 48 h post-transfection by PEG6000 precipitation.

### Quantitative RT-PCR

Endogenous target gene expression was analyzed following isolation of total RNA using the RNeasy Mini Kit (Qiagen, Hilden, Germany). First-strand cDNA was synthesized from 500 ng RNA using random hexamers and Verso cDNA Synthesis Kit (Thermo Fisher Scientific) according to the manufacturer’s instructions. qRT-PCR was performed using 1.5 μl of 1:5 diluted cDNA, 0.5 mM primers and DyNAmo ColorFlash SYBR Green qPCR Kit (Thermo Fisher Scientific) in 10 μl reaction volume using LightCycler 480 II instrument (Roche, Basel, Schweiz). Primer sequences used were as follows: *HPRT1*, forward: 5’-CCTGGCGTC ATTAGTGAT-3’, reverse: 5’-AGACGTTCAGTCCTGTCCAATAA-3’, *ALAS1*, forward: 5’-CTGCAAAGATCTGACCCCTC-3’, reverse: 5’-CCTCATCCACGAAGGTGATT-3’, *CEACAM6*, forward: 5’-CTCTACAAAGAGGTGGACAG-3’; *CEACAM6*, reverse: 5’-GTTAGAAGTGA GGCTGTGAG-3’; *Tns1*, forward: 5’-ACTACCTGCTGTTCAACCTC-3’, reverse: 5’-ATGA CAACTCCTATCCTGCC-3’, *Tns2*, forward: 5’-TATACTGCAAGGGAAACAAGG-3’, reverse: 5’-CTGCTGTTCATTCTGATGGAG-3’, *Tns3*, forward: 5’ CATTCATTGTTCGAGACA GCCA 3’, reverse: 5’-CAAATCTCCAGCTTTCTTGTTCAG-3’, *Tns4*, forward: 5’-GAGAGC AAGCAATCGAGCTG-3’, reverse: 5’-CAAAGTAGGGCTCCTCATCTG-3’, *Talin1*, forward: 5’-GAGTCAGTGTGCCAAGAACC-3’, reverse:5’-GTACACTTCTCCATTGTCTCCC-3’, *Vinculin*, forward: 5’-GATGAGGCTGAGGTCCGTAA-3’, reverse: 5’-AGCTCCTGCTGTCTCTCGTC-3’. Relative gene expression levels were calculated according to the 2^-ΔΔCT^ method. Results shown are averaged from at least three independent biological replicates.

### Immunofluorescence and Microscopy

For immunofluorescence analysis, 8x10^4^ cells (12-well plates) were grown on glass coverslips coated with type I collagen (0.5 mg/ml) or fibronectin (0.1 mg/ml), fixed with 3.7% formaldehyde in PBS for 15 min, and permeabilized with 0.2% (v/v) Triton X-100 in PBS for 10 min. For talin immunostaining, cells were fixed and permeabilized for 10 min in ice-cold (-20°C) Methanol/Aceton (1:1). After blocking in PBS, 10% FCS, 1% BSA, 0.05% Triton X-100 for 30 min, primary antibodies were diluted 1:100 to 1:400 and applied for 1 h at room temperature. Alexa-conjugated secondary antibodies were diluted 1:200 and applied for 1 h at room temperature. Samples were covered with ProLong Gold antifade reagent (Life Technologies) and imaged using a confocal microscope (TCS SP2 AOBS or SP5; Leica, Wetzlar, Germany) or an Apotome-containing Axio Observer.Z1 (Zeiss, Jena, Germany) equipped with 63x and 100x oil objectives and a monochrome Axiocam MRm camera. Representative images are shown.

For quantitative focal adhesion analysis, cells were allowed to adhere to collagen coated glass slides for 60 min and immunostained for paxillin. 43 single cells from n=4 experiments were recorded by a confocal microscope (SP5, Leica) with 100x oil objective. Paxillin fluorescent images were automatically batch processed and analyzed by the open source software ImageJ (version 1.48u). First, background fluorescence was removed by *substrate background* with a *100-pixel rolling ball radius*. Next, images were *thresholded* and binarized by the *Otsu-Method.* Finally, focal adhesion number and size area were analyzed above a lower threshold of 0.1 μm^2^. The aspect ratio was calculated as *major-axis/minor-axis* of focal adhesions *fitted ellipses*.

For 2D single cell area analysis, cells were seeded on collagen I or fibronectin-coated 8-well chamber slides (IBIDI). Cell attachment was allowed for 30 min before spreading was monitored over 15 h by time lapse analyses (10 min/frame) based on E2-Crimson red-fluorescence using a Leica SP5X inverse confocal microscope equipped with a Ludin cube life chamber, 40x oil objective and multi-positioning. Automated cell tracking was performed using the ‘CellMigrationAnalyzer’ tool of the MiToBo (http://www2.informatik.uni-halle.de/agprbio/mitobo/) package for ImageJ (http://imagej.nih.gov/ij/) to determine the mean area [μm^2^] of those single cells observed over a time period of at least 6 h. A total of 50−100 cells per condition were analyzed.

### Western blotting

Cells were washed twice with ice-cold phosphate-buffered saline and lysed in RIPA buffer (50 mM Tris/HCl, pH 7.4, 150 mM NaCl, 2 mM EDTA, 1% (v/v) Triton X-100, 0.1% SDS, Complete EDTA-free protease inhibitor cocktail) for 30 min on ice. The proteins were separated on SDS-PAGE, transferred to polyvinylidene fluoride (PVDF) membranes (Merck Millipore) and incubated with indicated primary antibodies used at 1:200 − 1:1000 dilution and incubated overnight at 4 °C. Fluorophore labelled secondary antibodies (LI-COR, Bad Homburg, Germany) were incubated for 1 h at room temperature. The fluorescence signals were detected with ODYSSEY CLx (LI-COR) and quantified by the associated software. Quantitative results were calculated from at least three independent biological experiments

### Active Rac1 quantification

Preparation of GST-PAK-CRIB fusion protein was carried out as described by Busche et al. (Busche et al., 2008). 4×10^6^ MDA-MB-468par or-susp cells were seeded in 15 cm dishes and cultured for 24 h. Afterwards, cells were lysed in 1 ml Rac lyses buffer (50 mM Tris pH 7.5, 1% Triton X-100, 150 mM NaCl, 10 mM MgCl_2,_ 0.5% sodium desoxycholate, 0.1% SDS, protease inhibitors). Equal amounts of protein lysates (1 mg) were incubated with immobilized GST-PAK-CRIB rotating at 4°C for 60 min and afterwards washed three times in Rac wash buffer (50 mM Tris pH 7.5, 1% Triton X-100, 150 mM NaCl, 10 mM MgCl_2_, protease inhibitors). Bound proteins were detected by western blotting using Rac1, GST and tubulin antibodies. Quantitative results of active pull down Rac1 levels were calculated and normalized to Rac1 in total lysates (normalized on tubulin).

### Cell Motility Assay

Migration assays were performed in transwell diffusion chambers (Corning Costar, Corning, NY, USA) with a pore size of 8 μm diameter as described previously (Wulfaenger et al., 2008). 600 μl RPMI1640 medium containing 10% FCS (optionally containing 50 ng/ml EGF) were added to the bottom of the 24-well plate. 4.5×10^4^ cells were inserted into the top chamber. After 22 h at 37°C, the upper side of the membrane was wiped to remove all remaining cells. The number of migrated cells was determined with the CellTiter-Glo Luminescent Cell Viability Assay (Promega, Madison, WI, USA) according to the manufacturer’s protocol. Luminescence was measured with a GloMax 96 Microplate Luminometer (Promega). Results are expressed as percentage of migrated cells compared to total cell number.

### Cell viability assay

2×10^4^ cells were seeded in a 96-well microplate for indicated time periods. Proliferation was determined with Colorimetric Cell Viability Kit I (PromoKine, Heidelberg, Germany) according to the manufacturer’s protocol. Absorbance at 450 nm was measured after 1.5 h of incubation at 37°C using a microplate reader.

### Apoptosis assay

1×10^6^ cells were subjected to the EnzChek Caspase-3 Assay Kit #2 (Thermo Fisher Scientific) according to the manufacturer’s protocol. Absorbance at 520 nm was measured after 30 min of incubation at room temperature using a microplate reader.

### Adhesion assay

For cell-substratum adhesion assays, 96-well tissue culture plates were coated with type I collagen (0.5 mg/ml) or fibronectin (0.1 mg/ml) in PBS at RT overnight, rinsed twice with PBS and cells were allowed to adhere to the coated surface for 30 min and 60 min respectively. To remove non-adherent cells, wells were washed twice with PBS and DNA was stained with DAPI. Fluorescence was measured with ClarioStar High Performance Monochromator Multimode Microplate Reader (BMG LABTECH, Ortenberg, Germany).

### Collagen contraction assay

Increasing cell numbers (1×10^4^, 3×10^4^, 5×10^4^ and 10×10^4^) of MDA-MB-468par and-susp cells were mixed with type I collagen (final concentration 1.5 mg/ml) in RPMI1640 medium containing 10% FCS and 10 ng/ml EGF and allowed to solidify in 96-well plates. For the negative control collagen I was mixed only with the according medium. After 1 h incubation at 37°C 200 μl RPMI1640 medium containing 10% FCS and 10 ng/ml EGF were added to each well. The collagen lattices were detached from the surface of the well and after 4 days incubation at 37°C the total area covered by the collagen lattices was measured and compared to the negative control. Results show the shrinkage of the collagen gel in %.

### Flow cytometry

5×10^5^ cells were centrifuged at 300xg for 10 min. Cells were resuspended in 100 μl PBS (pH7.2 containing 0.5% BSA and 2mM EDTA) and incubated for 10 min at 4°C with 5 μl anti-integrinß1, ß3, ß4 and α2, α5, α6, αV antibodies conjugated with FITC, PE and APC. Cells were washed by adding 1 ml of PBS and resuspended in 100 μl PBS. Fluorescence was measured with BD Acccuri Flow Cytometer. Integrin surface levels in 468susp cells were compared to 468par cells.

## Acknowledgements

We thank Nadine Bley and the Core Facility Imaging for assistance with microscopy and adhesion assays, Stefan Hüttelmaier for kindly providing the vector pMD2.G, psPAX2 and pLVX-shRNA2, Reinhard Hoffmann for the transcriptome data analysis, Elisabeth Bormann for technical help with Figure 7, and Pascal Rudewig and Anja Weber for excellent technical assistance.

## Author contributions

AV and UB performed experiments, analysed the data and wrote the manuscript, LL selected cell lines, performed the microarray, and analysed the data, AK and AE performed experiments, MS conceived the project and designed experimental approaches, GP conceived and supervised the project, analysed the data, and wrote the manuscript.

## Figures legends

***Supplementary Figure S1. Characterization of 468^par^ and 468^susp^ subpopulations***. The mRNA expression levels of Tns1, Tns2 and Tns4 (**A**), Vinculin (**B**) and Talin (**C**) were determined by qRT-PCR normalized to ALAS1 and HPRT1 in 468^par^ and 468^susp^ cells. All error bars represent ± SEM, n≥3. All p-values were calculated using an unpaired one-sample student’s t-test (*p≤0.05, ** p≤0.01).

***Supplementary Figure S2. Adhesion dependent expression of Tns3 in MCF7 cells.***

**(A)** MCF7 cells were cultivated on polyHEMA coated dishes for up to 72 h. Afterwards cells were allowed to readhere to normal culture dishes. Scale bars, 50 μm **(B)** Tns3 mRNA transcript levels from indicated conditions were determined by qRT-PCR and normalized to ALAS1 and HPRT1. **(C)** Quantification of Tns3 protein levels in total cell lysates from indicated conditions normalized to tubulin. **(D)** Quantification of vinculin protein levels in total cell lysates from indicated cells normalized to tubulin. **(E)** Quantification of talin protein levels in total cell lysates from indicated cells normalized to tubulin. All error bars represent ± SEM, n=3. P-values were calculated by Tukey’s post-hoc test using one-way ANOVA (* p≤0.05, *** p≤0.001).

***Supplementary Figure S3. Impaired adhesion of HCT8 colon cancer cells depleted of Tns3.* (A)** Tns3 levels in parental HCT8 cells, and HCT8 cell stably transfected with control or Tns3-specific shRNA. Representative western blots of total cell lysates. **(B)** Adhesion of stable HCT8 cell lines to collagen-coated surfaces for 60 min. Shown is the percentage of adhered cells normalized to the control. Error bars represent ± SEM, n=3. * p≤0.05 according to unpaired one-sample student’s t-test.

***Supplementary Figure S4. Tns3 knockdown in MDA-MB-468 using specific siRNAs***. **(A)** Phase contrast images of 468 cells transfected with different Tns3 specific siRNAs and ctrl siRNA respectively. Cells were additionally stained for F-actin. Scale bars are 50 μm and 10 μm, respectively. **(B)** Western blot of Tns3 in total cell lysates from indicated cells. Tubulin was used as a loading control.

***Supplementary Figure S5. CEACAM6 and E-cadherin in MDA-MB-468 subpopulations***. **(A)** Transcriptome profiling of the 468^par^ cells (3 independent RNA preparations) and 468^susp^ cells (RNA preparations of 3 independent selections) using the Human Gene ST array (Affymetrix) showed differentially expressed CEACAM6. Depicted are mean microarray signal intensities. **(B)** CEACAM6 mRNA transcript levels were determined by qRT-PCR normalized to ALAS1 and HPRT in 468^par^ and 468^susp^ cells. Error bars represent ± SEM, * p≤0.05 according to unpaired one-sample student’s t-test (n=3). **(C)** Western blot of E-cadherin in total cell lysates from indicated cells. Tubulin was used as a loading control. **(D)** Representative images of indicated cells plated on glass coverslips before fixation and stained for E-Cadherin (green) and F-actin. Merged images show E-Cadherin (green), F-actin (red) and DNA (blue). Scale bars, 20 μm.

